# Genomic evidence supports the introgression between two sympatric stickleback species inhabiting the White Sea basin

**DOI:** 10.1101/2020.11.24.396010

**Authors:** Artem Nedoluzhko, Fedor Sharko, Svetlana Tsygankova, Eugenia Boulygina, Amina Ibragimova, Anton Teslyuk, Jorge Galindo-Villegas, Sergey Rastorguev

## Abstract

Interspecies hybridization is driven by a complex interplay of factors where introgression plays an important role. In the present study, the transfer of genetic material, between two quite distant fish species from different genera, through spontaneous hybridization was documented with dedicated molecular and bioinformatics tools. We investigate the genomic landscape of putative stickleback-relative introgression by carefully analyzing the tractable transposable elements (TE) on the admixed genome of some individuals of two sympatric stickleback species inhabiting northwestern Russia, namely the three-spined (*Gasterosteus aculeatus*) and the nine-spined (*Pungitius pungitius*) sticklebacks. Our data revealed that unique TE amplification types exist, supporting our proposed hypothesis that infers on the interspecific introgression. By running a restriction site-associated DNA sequencing (RAD-Seq) with eight samples of *G. aculeatus* and *P. pungitius* and subjecting further the results to a contrasting analysis by variated bioinformatic tools, we identified the related introgression-linked markers. The admixture nature observed in a single sample of the nine-spined stickleback demonstrated the possible traces of remote introgression between these two species. Our work reveals the potential that introgression has on providing particular variants at a high-frequency speed while linking blocks of sequence with multiple functional mutations. However, even though our results are of significant interest, an increased number of samples displaying the introgression are required to further ascertain our conclusions.

## Introduction

Interspecific hybridization in animals has been considered for a long time as an abnormal process leading to the destruction of reproductive isolation [1]. In the early 20th century, supporters of this viewpoint increased after the description of postzygotic isolation (PSI) [2]. The PSI supposes that the allele incompatibility increases as the square of the genetic distance between species complicate the hybridization between distant species. Nevertheless, while the interspecific hybridization is a natural event, it is also an active participant in some evolutionary process as the speciation or the new trait acquisition. In support of the same, it has been demonstrated that 25% of the plant species and more than 10% of animals from different taxa present traces of hybridization [3,4]. In contrast, the theoretical foundations of hybridogenic speciation propose the formation of reproductive barriers between hybrids and parents as the possible genetic mechanisms [5]. Fishes are not the exception, and thus the interspecific hybridization events have been appreciated as a quite common event among this phylogenetic group [6,7].□

Nevertheless, the following logical question is placed forefront among the previously described concepts. Why, despite the exitance of PSI, hybridization is still a present common feature in nature? In this regard, variated hypothesizes attempt to address the critical role of the hybridization in evolution. While still in debate, the most sounded hypothesis establishes a dramatic acceleration of the natural selection power by increasing the genetic variability resulting from the same [8–10]. However, studies demonstrate that hybridization allows offspring to obtain new traits [11–13], and conquer new habitats and ecological niches [8–10,14]. Such studies showed that hybridization is also part of the vertebrate speciation [5,15].

The high-throughput sequencing and genotyping era opened great avenues for studying in detail the hybridization involvement in the evolutionary processes [16]. To do it so, the species from the family Gasterosteidae (order *Gasterosteiformes*) are accessible elements to perform evolutionary studies, speciation, and several aspects of functional biology [17–20]. Notably, the three-spined stickleback (*Gasterosteus aculeatus*) is a well-studied model vertebrate species that acquired the freshwater adaptation due to the “freshwater adaptive standing genetic variation” – the quite extended genomic loci that significantly differ between marine and freshwater forms of this species [21,22]. The marine three-spined stickleback ancestor perhaps obtained these “divergence islands” from a close related freshwater species. Just recently, the hybridization between the marine and freshwater forms of the three-spined stickleback has been proved by using genetic and morphological data. Besides, an extended ecological potential in the hybrid offspring has been clearly described when comparing the hybrids with the parent forms [23–25]. Conversely, it is well recognized that some species from the genus *Pungitius*, like *P. pungitius* with *P. tymensis*, or *P. platygaster* with *P. pungitius* can form fertile interspecific hybrids with the ability to produce viable backcrosses [26,27]. However, according to recent literature, intergeneric hybrids of sticklebacks have not been described so far.

In the present research, we hypothesized the possible genomic introgression between two separate Gasterosteidae species belonging to two distinct genera, *Pungitius* and *Gasterosteus*. To validate our hypothesis, we first ascertain through the exhaustive analysis of several molecular pieces of evidence achieved by carefully analyzing of transposable elements (TE) and restriction site-associated DNA sequencing (RAD-seq) on a small number of specimens. Subsequently, the application of different bioinformatics approaches provided novel interspecies hybridization marks specific for the Gasterosteidae. Besides, our results also put forward a solid foundation on the use of transposable element analysis as a novel, low time consuming, and affordable method within several population-wide studies designed to find intergeneric genomic introgression [28]. In the meanwhile, we predict that further population genome-wide studies will shed light on this new hypothesis.

## Results

A TE analysis was performed to determine the possible presence of introgressive hybridization between the three- and nine-spined stickleback species. DNA electrophoresis of the TE PCR products revealed that various types of elements are differentially amplified among the analyzed samples. However, in two loci, the Ltr89 and Tcl5, the TE were equally represented in both species' samples. For all the remaining loci, only the three-spined stickleback samples TE were mostly constant (**Figure 1**).

**Figure 1.**
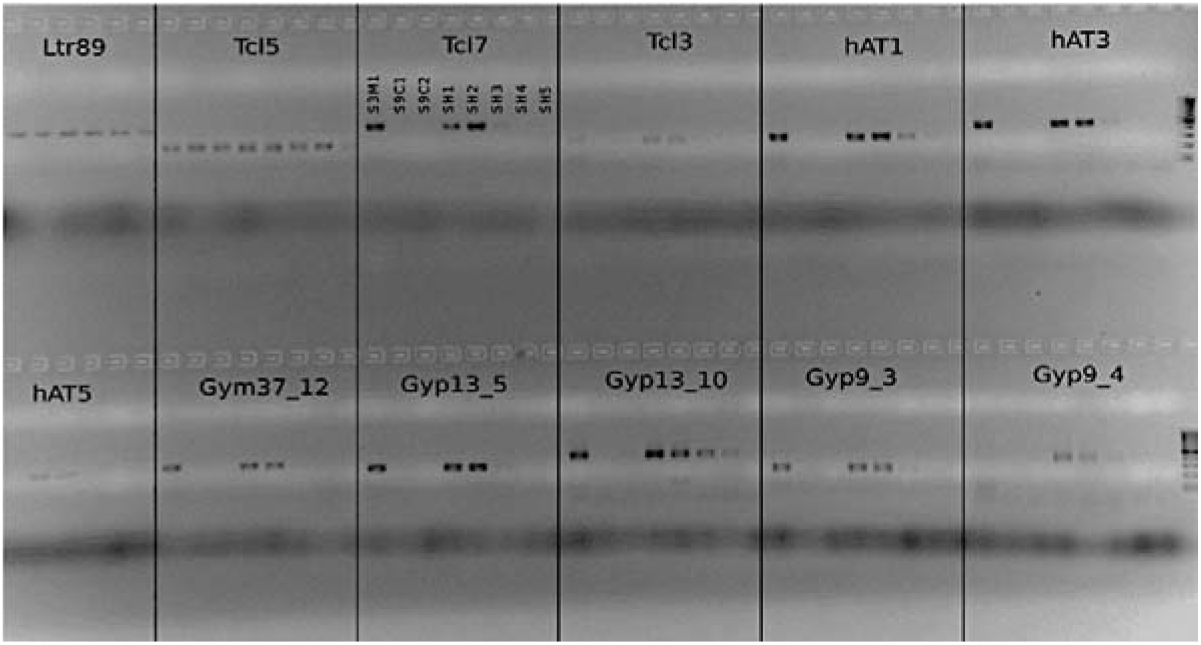
Gel electrophoresis confirming the presence of three-spined transposable elements (TE) in the nine-spined stickleback samples. To avoid overwriting each sample name and order is only indicated in the Tcl7 locus, but the same applies to each of the 12 loci presented.

Strikingly, the SH3 sample, suspected as an admixed specimen of the nine-spined stickleback, produced a clear noticeable signal in most amplified profiles. Nevertheless, weaker than in the three-spined fish samples. Besides, some remaining samples of the nine-spined fish either show a weak signal at several loci (e.g., gyp13_10). Whatever the case, the fact that some nine-spined samples showed the presence of the three-spined like TE provided the required evidence for conducting further analysis focused on expanding the genomic introgression knowledge between the three- and nine-spined sticklebacks.

To gain further insights into the genomic signals of introgression in the Gasterosteidae family, using the Illumina platform, we performed a RAD-Seq analysis of the DNA isolated from the eight stickleback samples described below (see: **Table 5**). The number of Illumina reads produced and mapped for all samples are shown (**Table 1**).

**Table 1.**
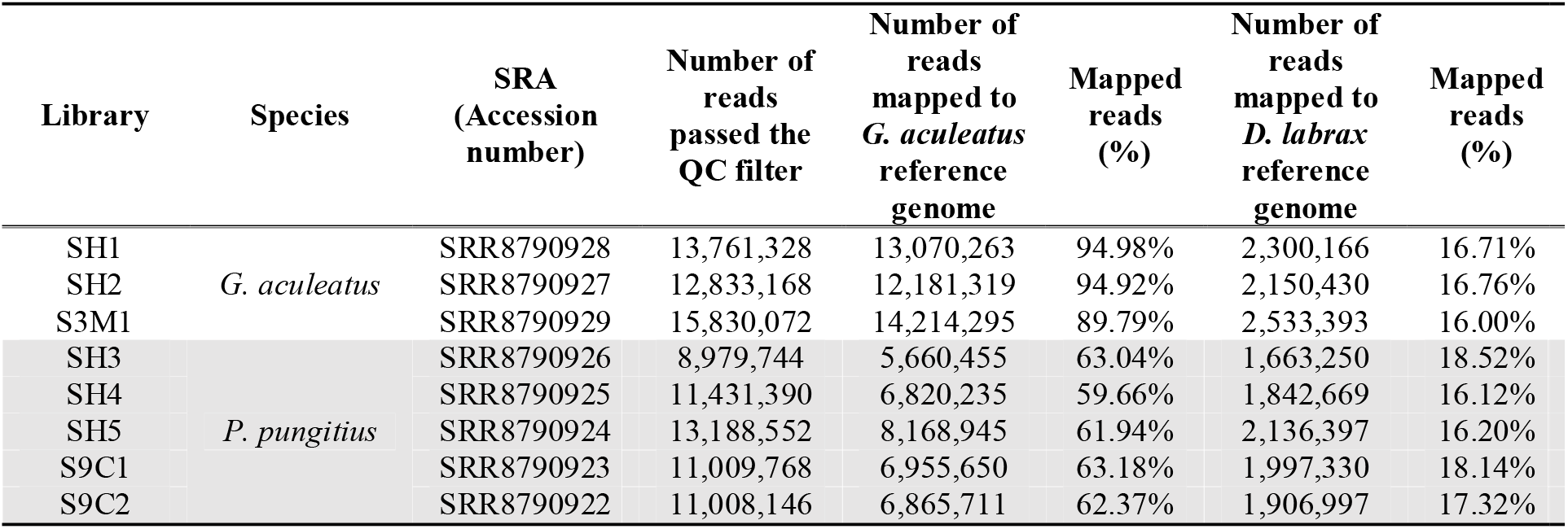
Illumina generated reads and mapping statistics on obtained reads mapped to three-spined stickleback and European sea bass genomes.

Resulting from mapping the RAD-Seq data to the dicLab v1.0c reference genome, 108,969 loci in all samples were determined. However, after converting the vcf files to the genlight (adegenet) format, the total loci number decreased by 4,062. The reduction observed in the loci number resulting from the conversion is attributed to the particular characteristic of the genlight format that does not support loci with more than two alleles. We then created a distance matrix for all samples based on their genotypes' dissimilarity and conducted a cluster analysis based on the Neighbor-Joining method (**Figure 2**).

**Figure 2.**
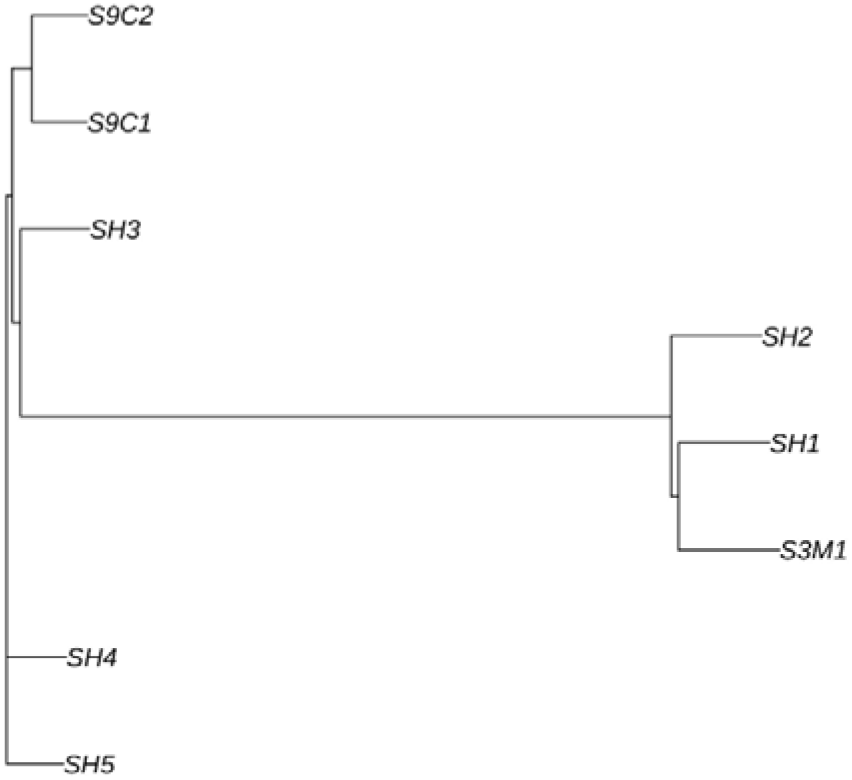
Cluster analysis for the three- and nine-spined stickleback samples, based on Euclidean distances between their genotypes.

The genotypes defined in the vcf file were used for the admixture analysis in the Structure tool [37]. The Structure analysis of the nine-spined stickleback SH3 sample confirmed our previous assumption of this specimen's admixture origin. Besides, the SH2 sample of the three-spined stickleback specimen, collected in the same location as the SH3 sample, also displayed traces of intergeneric hybridization in its genome. However, despite the source, the admixture value present in both cases is less than one percent, which is extremely low but in a large loci number, supporting the reliability of the present result.

Based on the structure analysis, the inferred ancestry of individuals in the SH2 and SH3 libraries obtained from the three- and nine-spined sticklebacks, respectively share contrasting clusters on the 0.7 % level (**Table 2**). The quantitative values of an admixture could be revised because only a part of the whole genomes was analyzed. Moreover, we used only the available reference genomes from distant species, but the possibility of intergeneric genomic introgression is clearly inferred.

**Table 2.** Inferred ancestry of individuals based on the Structure analysis

| Library | Species | Inferred clusters |  | 90% probability interval |  |
| --- | --- | --- | --- | --- | --- |
| SH1 | *G. aculeatus* | 1.000 | 0.000 | 0.999,1.000 | 0.000,0.001 |
| SH2 |  | 0.993 | 0.007 | 0.991,0.995 | 0.005,0.009 |
| S3M1 |  | 1.000 | 0.000 | 0.999,1.000 | 0.000,0.001 |
| SH3 | *P. pungitius* | 0.007 | 0.993 | 0.005,0.008 | 0.992,0.995 |
| SH4 |  | 0.000 | 1.000 | 0.000,0.001 | 0.999,1.000 |
| SH5 |  | 0.000 | 1.000 | 0.000,0.001 | 0.999,1.000 |
| S9C1 |  | 0.000 | 1.000 | 0.000,0.001 | 0.999,1.000 |
| S9C2 |  | 0.000 | 1.000 | 0.000,0.001 | 0.999,1.000 |

We selected three-spined - S3M1 and nine-spined - S9C1 and S9C2 specimens as test samples, and all other nine-spined specimens compared iteratively against them. D-statistics value increased for SH3 sample (**Table 3**). The value of 0.32 and 0.34 for S9C1 and S9C2 respectively are positive, which means there was an allelic frequency shift in the direction of convergence of the nine-spined SH3 sample with an S3M1 three-spined sample. Z-score of 3.9 means that the confidence of the admixture is high [40]. The same results we obtained when other three-spined and nine-spined specimens, accordingly, were used as test samples (data not shown).

**Table 3.** Results of genomic introgression test using the European sea bass (*Dicentrarchus labrax*) as outgroup.

| Outgroup | Population1 | Population2 | Population3 | D-stat | Z-score | ABBA | BABA | Number of SNPs |
| --- | --- | --- | --- | --- | --- | --- | --- | --- |
| *D.labrax* | S3M1 | S9C1 | SH3 | 0.3258 | 3.899 | 420 | 220 | 10,044 |
| *D.labrax* | S3M1 | S9C1 | SH4 | 0.0223 | 0.276 | 270 | 260 | 10,044 |
| *D.labrax* | S3M1 | S9C1 | SH5 | 0.0774 | 0.8 | 280 | 240 | 10,044 |
| *D.labrax* | S3M1 | S9C2 | SH3 | 0.346 | 4.916 | 440 | 210 | 10,044 |
| *D.labrax* | S3M1 | S9C2 | SH4 | 0.0542 | 0.707 | 260 | 230 | 10,044 |
| *D.labrax* | S3M1 | S9C2 | SH5 | 0.1137 | 1.251 | 280 | 220 | 10,044 |

We conducted an additional introgression test based on ancestry graph models, implemented in the TreeMix software package by using a large number of SNPs to estimate the historical relationships among populations. The migration events in the TreeMix test splits and goes from the root node of the three-spined subgraph to the nine-spined sample (SH3) as well as from the nine-spined subgraph to the three-spined sample (SH2) (**Figure 3**). Simultaneously, the latter result with SH2 specimen was not supported by Structure and D-statistics analyses.

**Figure 3.**
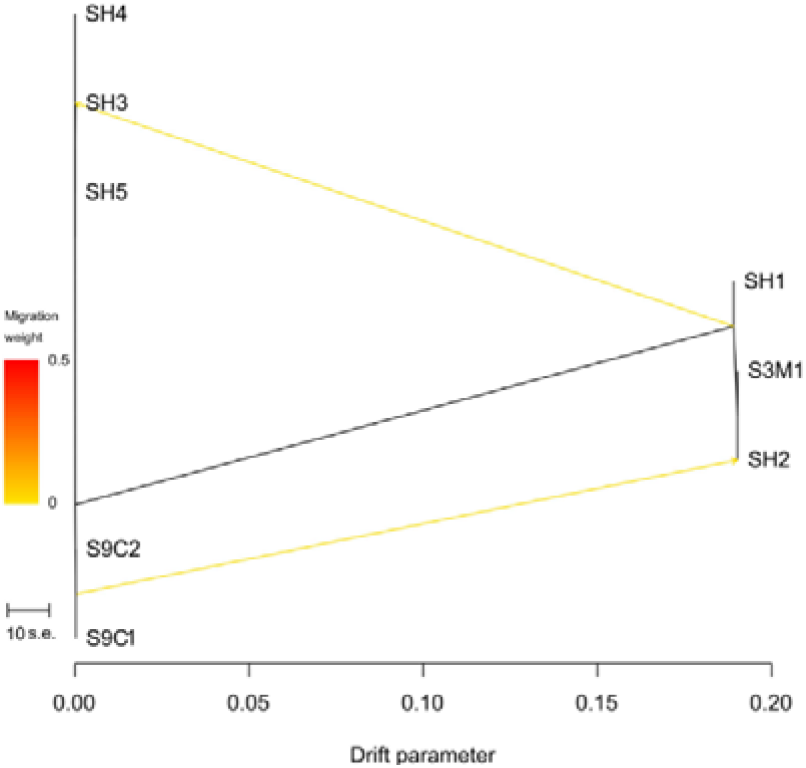
The evidence for admixture in the two stickleback species obtained by the TreeMix software analysis.

Despite that each specimen was collected, carefully stored on individual tubes, and followed a strict procedure on processing the DNA library, a remote possibility of DNA contamination among samples of the three- and nine-spined specimens exist. At first glance, this fact could explain the unexpected admixture results. To exclude the possibility that a sample has been mixed and sequenced, an assessment of mapped data for mismatches was conducted. The results revealed that SH3 admixed sample does not differ significantly from the rest of the samples. Even more, the SH3 admixture has the smallest mismatch rate comparing other specimens (**Table 4**).

**Table 4.** Test for contamination. Relative mismatch quantity in mapping data of three-spined and nine-spined stickleback specimens.

| Specimen | S3M1 | S9C1 | S9C2 | SH1 | SH2 | SH3 | SH4 | SH5 |
| --- | --- | --- | --- | --- | --- | --- | --- | --- |
| Mismatch quantity | 0.07941 | 0.07559 | 0.07548 | 0.08059 | 0.07905 | 0.7356 | 0.07691 | 0.07621 |

Therefore, these results expressing the relative mismatch quantity in mapping data, sufficiently support the exclusion on the possible admixed sample contamination.

## Discussion

Hybridization between evolutionary distant animal species has been traditionally perceived as a rare event in nature [43–45]. However, recent studies propose the introgressive hybridization as a widespread phenomenon among closely related vertebrate species [46,47]. Interestingly, this feature is particularly frequent among teleost fish [48,49]. Nevertheless, this effect is quite probably related only due to the total number of fishes exceeding by far that of all other vertebrates. Whatever the case, the new insights gained from different models and non-models teleost species and sequencing projects have recently revealed several peculiar features of fish genomes that might have played a particular role in fish evolution and speciation [50].

In this study, our efforts were focused on providing a multilevel survey on the evaluation of a possible introgression between the three-spined and nine-spined stickleback species. For this purpose, different powerful bioinformatical methods which have shown previous success on linking loci and correlating allele frequencies [37], revealing admixture processes [40], and inferring population splits and mixtures from genome-wide allele frequency data [41], were approached.

Besides, to increase the data accuracy, we developed a TE PCR-based identification system that allows for finding traces of previous introgression events between diverse stickleback species. The introgression process usually induces the burst of TE in the resulting hybrids [28]. Therefore, it is assumed that TE play essential roles in animal evolution, suggesting this part of the genome as a very sensitive marker of introgression. In our study, resulting from the emerged TE PCR products, and following an electrophoretic approach, we gave light on the apparent variation on the differential amplification of the TE between the two distant sympatric species of *Gasterosteiformes*, the three- and nine-spined sticklebacks.

The interspecies hybridization was expected to increase the TE activity in their hybrid offspring. In agreement with our findings, it has been previously shown that interspecific hybridization disrupts the stability of this part of the genome resulting in mutations and genetic instability [51]. We found that a single nine-spined (SH3) stickleback sample, presenting a reduction in the number of spines, showed TE specific to the three-spined stickleback (particularly, Tcl7, hAT1, hAT3, and Gyp9_4). The emerging evidence revealed that our novel approach succeeds in providing the required evidence on TE-elements' usage as an excellent molecular marker for introgression detection in fish. Thus, as a highlight of the present study, we introduce a novel TE PCR approach that provides fast and accurate evidence of the admixture process between species from the different genera - *G. aculeatus* and *P. pungitius*.

RAD-Seq has been proposed as the tool of choice in introgressive hybridization studies across individuals of non-model organisms [52,53]. For example, using RAD-Seq a secondary hybridization event at the end of the last glaciation period between two sole African species, *Solea senegalensis*, and *S. aegyptiaca*, has been reported despite a lengthily allopatric separation exists [54]. Furthermore, using the same tool, the genetic admixture and interspecies hybridization between different cichlid species from the *Alcolapia* genus inhabiting the African lake Natron has been recorded [55]. Together, these examples emphasize the importance of the RAD-seq analysis for interpreting hybridization patterns. In the present study, the RAD-Seq analysis revealed a strong relationship among the adaptive allelic character sets of the freshwater nine-spined (*P. pungitius*) stickleback acquired from its three-spined marine relative during the introgression process.

Note that here we did not use a well-annotated *G. aculeatus* genome. Instead, the European sea bass genome sequence served as reference, avoiding in this way the possible bias in the mapping efficiency. Besides, several approaches, including an assessment of mapped data for mismatches, and contamination test, were followed to increase the confidence in the results obtained from a limited number of specimens in this proof-of-concept investigation. So, thus far, the comparative bioinformatical analysis based on Structure, Treemix, and D-statistics methods of genomic data for this and other three-spined and nine-spined stickleback samples provides a definite milestone on the suspected admixture origin of SH3 specimen. The possibility of such hybridization events was explored toward the interspecific level [56]. However, only a few but sporadic findings showed the examples of intergeneric hybridization [49,57]. Thus, our results suggested that introgressive hybridization between the three- and nine-spined stickleback species adds a new case example of intergeneric hybridization.

Besides, we inferred that the newly TE-based PCR analysis should become a convenient type of preliminary experiment in case of in-depth screening of the populations where interspecies and intergeneric hybridization are expected. Whatever the case, we anticipate that further population-wide genomic studies will shed light on the adaptivity hypothesis of intergeneric hybridization between Gasterosteiformes.

Interestingly, along with the molecular findings, the ecological characteristics of the inhabiting aquatic niches of both stickleback species either implies a strong challenge to overcome. While the three-spined lives in a gradient between the marine and freshwater environments, the nine-spined stickleback is predominantly a freshwater inhabitant [18]. Previous studies in teleost fish have found that sensing osmotic stress by the skin cells through a transient potential receptor vanilloid 4, the differential activation or repression of several genes related to variated physiological functions could be induced [58,59]. Indeed, our study's particular relevance is that both admixture samples have been sampled in the stream's estuary, where both species coexist, and environmental changes are more pronounced due to the tidal activity. However, the three-spined (*G. aculeatus*) has a higher tolerance of salinity than the nine-spined (*P. pungitius*). Recently, a transgenerational plasticity process based on epigenetic marks and epimutations has been proposed to contribute to adaptation and acclimation of stickleback to salinity [60]. However, despite the exciting findings reported, such epigenetic studies and our genomic data interplay still maintain the salinity adaptation capacity of *G. aculeatus* unresolved. Thus, subsequent studies showing if *P. pungitius* requires the translocation of critical alleles of *G. aculeatus* to colonize into the seawater environment are guaranteed.

## Conclusions

In this paper by using several independent methods, we managed to provide a hypothesis on the possibility of genomic introgression between two genera of sticklebacks (*Pungitius* and *Gasterosteus*) in northwestern Russia. Besides, a new molecular method with high power and few false positives over other related tests was provided. This convenient TE PCR marker system, developed, may contribute to fast and quality sampling of the specimens in the upcoming population-wide studies of Gasterosteiformes, with intergeneric hybridization traces. In short, if the primer system amplifies *Pungitius* specimens, the probability of *Gasterosteus* introgression would be high in that sample. We pre-accept that this method is sensitive to the group genomic introgression due to TE copies' multiplicity in the genome of animals.

## Materials and Methods

### Fish Sampling

Two sympatric species of the Gasterosteidae family commonly inhabiting variated niches in the Northern hemisphere, namely the three-spined (*Gasterosteus aculeatus*), and the nine-spined (*Pungitius pungitius*) stickleback were captured in the coastal lands of Chkalovsky village in the Republic of Karelia, Russia (**Table 5**).

**Table 5.**
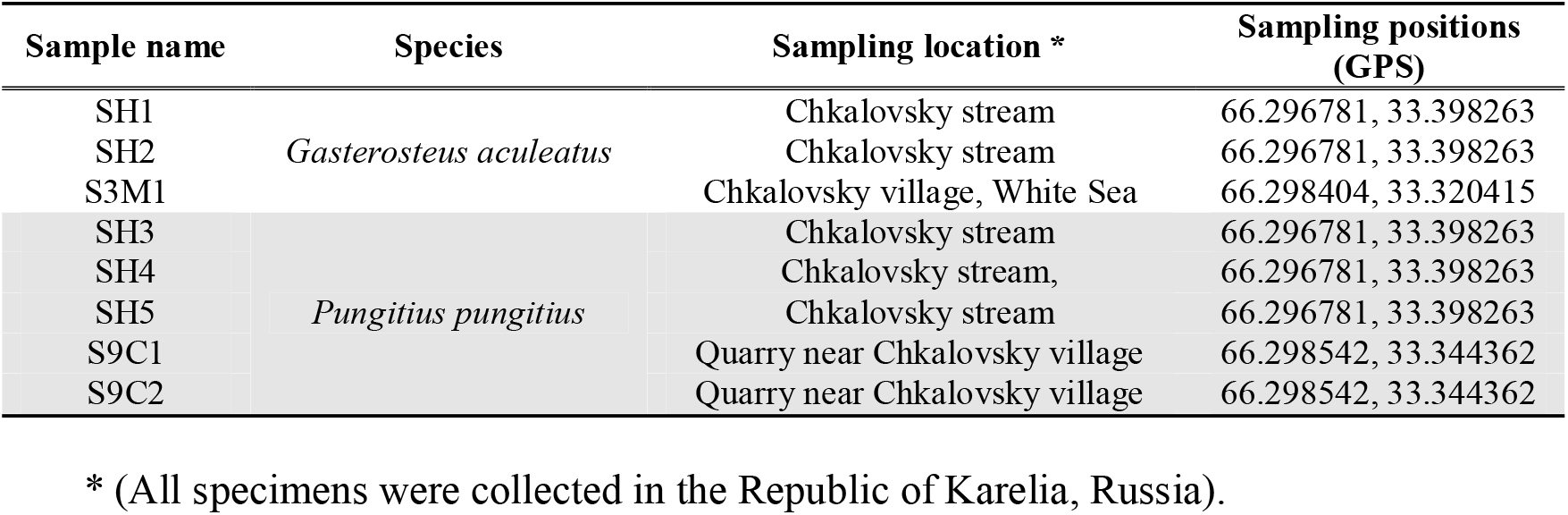
Nomenclature and sampling location of the three- and nine-spined stickleback samples were used on this study's variated analyses.

Individuals of both species were collected across a salinity gradient from the brackish White Sea waters to the freshwater in the stream within the same geographical area. Indeed, three-spined stickleback samples were collected from the White sea, near the Chkalovsky village (**Figure 4-A**). Samples of both species were obtained from the freshwater stream in a portion flowing into the White sea (near the tidal zone), where both species cohabitate (**Figure 4-B**). Nine-spined samples were collected from the freshwater quarry (**Figure 4-C**).

**Figure 4.**
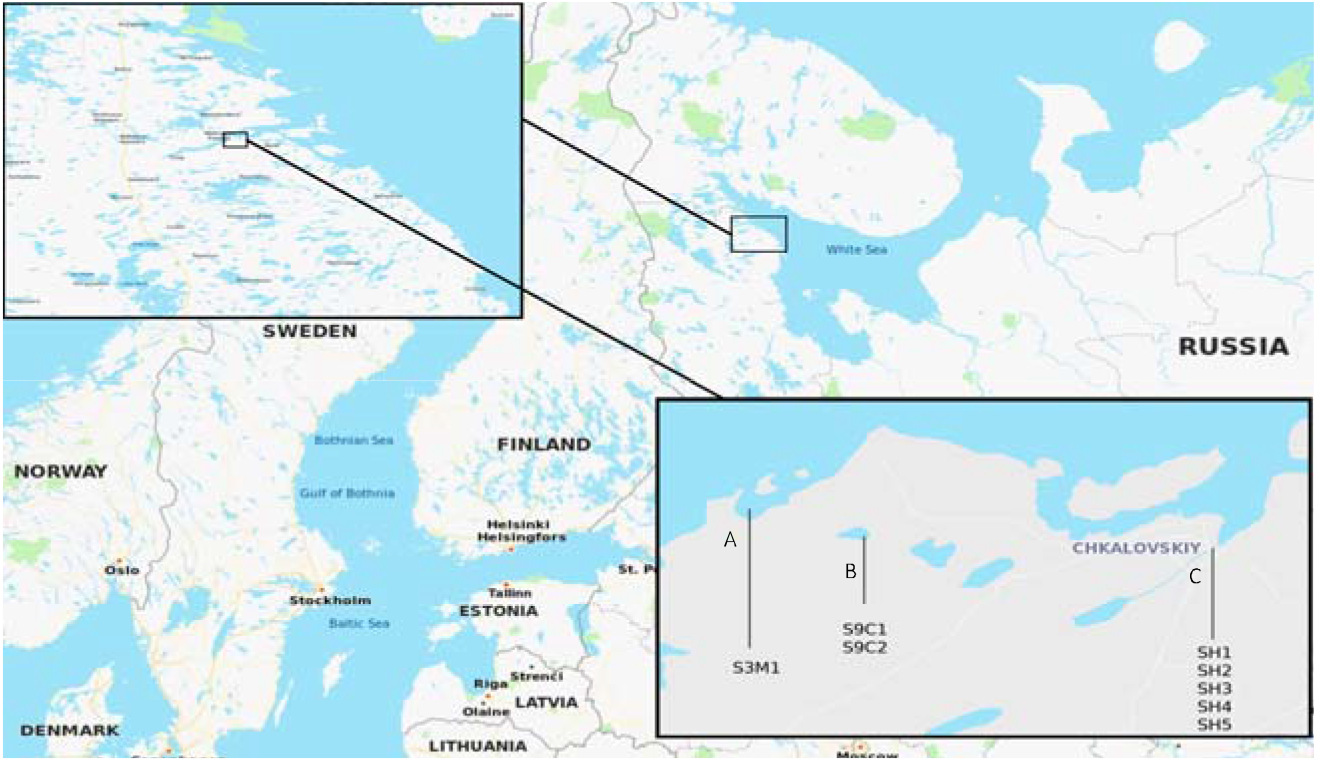
Map showing the different locations in Karelia, Russia, where *Gasterosteus aculeatus* (Three-spined) and *Pungitius pungitius* (Nine-spined) stickleback samples were collected. **A**) Chkalovsky village (S3M1), **B**) Chkalovsky quarry (S9C1 and S9C 2), **C**) Chkalovsky stream (SH1 to SH5). (See Table 1 for further details).

After phenotypical validation of each stickleback specimen, conducted by *in situ* dorsal fin spine counting, the anal fin was clipped and preserved in 70% ethanol for genomic DNA isolation, as described previously [29]. All specimens were caught and immediately released without any further morphological analysis to avoid stressing much the fish. One of the nine-spined sticklebacks (SH3 sample) presented a reduction in the number of spines, yet it was also used to conduct genetic analysis. This work was carried out in accordance with relevant guidelines and regulations and was approved by ethical committee of Institute of Bioengineering, Research Center of Biotechnology of the Russian Academy of Sciences, Moscow, Russia.

### DNA extraction, and Transposable Element (TE) Analysis

Total DNA was extracted from each stickleback sample by using the routine phenol-chloroform method. Each DNA sample concentration was quantified using a Qubit fluorimeter (Thermo Fisher Scientific, USA). The TE analysis is one of the most sensitive methods for introgression detection in animals [30]. In this work, the publicly available published sequences of the three-spine stickleback TE have been used [31]. Besides, we designed specific oligonucleotide PCR primers for amplifying variated TE fragments of several stickleback samples available. Illumina nucleotide reads of the Japanese nine-spined stickleback available at the NCBI sequence read archive (DRX012173) were used to design the specific primers for TE PCR. The Japanese nine-spined stickleback reads were mapped to the three-spined stickleback TE as the reference. Primers were designed to amplify TE's fragments that are not covered by Japanese nine-spined stickleback DNA reads. Thus, these primers could amplify only the three-spined stickleback TE but did not do so on the nine-spined ones. As a consequence, twelve TE were selected, five RNA, and seven DNA transposons, respectively. The resulting TE and the oligonucleotide primer sequences are presented (**Table 6**).

**Table 6.**
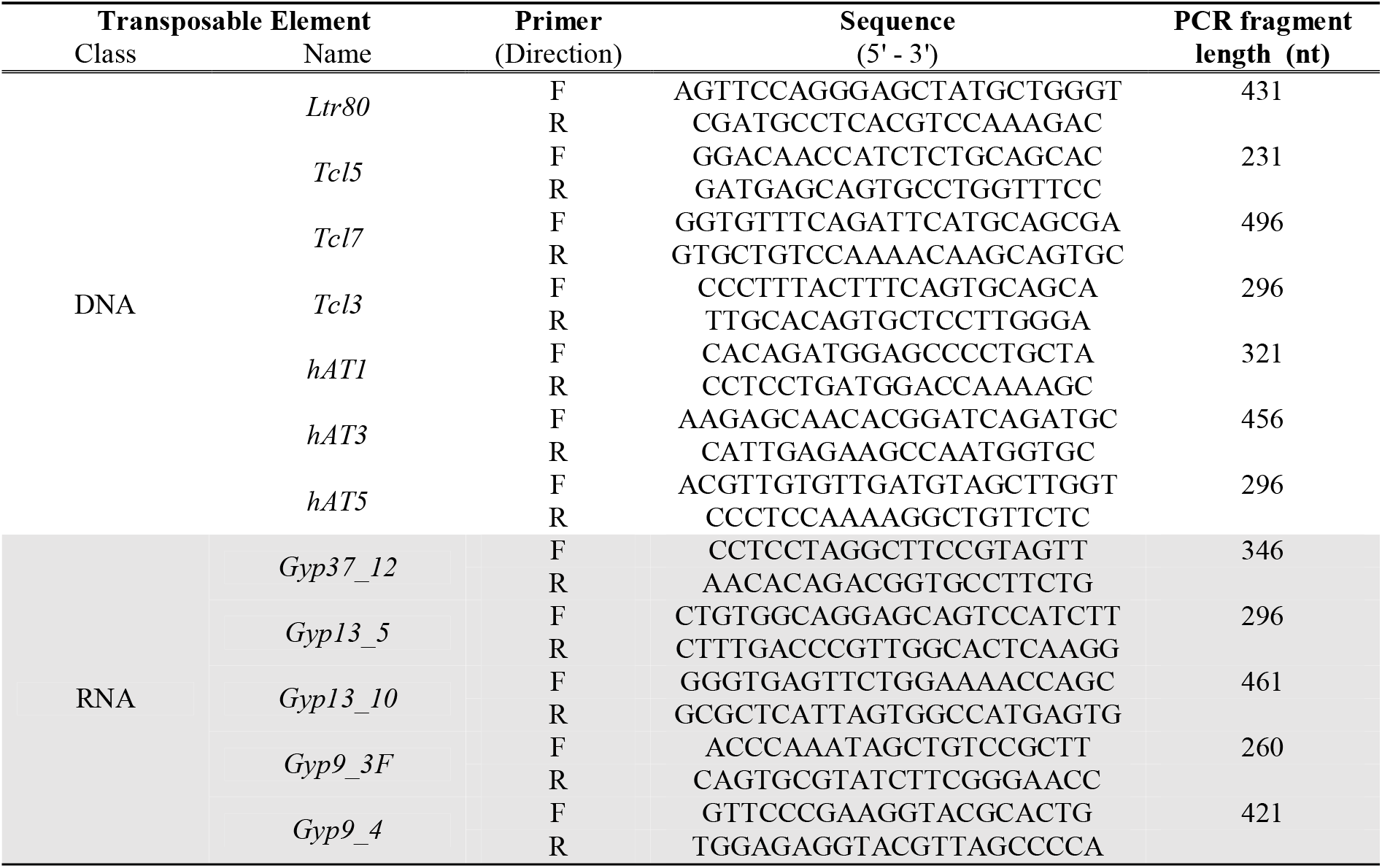
Transposable element (TE) of *G. aculeatus* and the primer list used for their amplification.

Encyclo PCR Kit (Evrogen, Russia) was used for the evaluation of the transposable element analysis. Same conditions for all loci PCR amplification were used: initial denaturation of 95°C for 10 minutes followed by 35 cycles of 94°C for 20 sec., 58°C for 30 sec.; 72°C for 60 sec.; and final extension for 10 min at 72°C. The PCR amplification products were separated by electrophoresis in 1% agarose gel with ethidium bromide. Universal Hood II Gel Doc System (Bio-Rad Laboratories, USA) was used for the electrophoresis visualization.

### Library preparation, and RAD-sequencing (RAD-Seq)

The genomic DNA was digested by using two nucleases, the *EcoRI*, and *MspI* (New England Biolabs, UK). The fragmentation estimation and the size-selection were carried out using 1% agarose gel electrophoresis. DNA fragments between 350 and 450 bp were selected and extracted from the agarose gel. The obtained DNA was purified using the QIAquick Gel Extraction Kit (QIAGEN, USA). Approximately 1 µg of fragmented DNA was used for each library preparation using the NEBNext® DNA Library Prep kit (New England Biolabs, UK); DNA-libraries were multiplexed with NEBNext Multiplex Oligos for Illumina kit (New England Biolabs, UK). According to standard Illumina cluster generation and sequencing protocols, we sequenced the libraries in an Illumina 2500 platform (Illumina, USA). On this basis, 100-bp paired-end reads were generated. The data were submitted to the Sequence Read Archive under project number PRJNA529064.

### Illumina Data Analysis

The DNA reads were mapped only to reference genome of *G. aculeatus* (BROAD S1, Ensemble database version 95.1) because the nine-spined stickleback genome is currently unavailable. Mapping on the three-spined stickleback genome showed a bias in the mapping efficiency between three- and nine-spined samples RAD-Seq data, that may lead to undesirable effects in the data analysis. Therefore, the European sea bass (*Dicentrarchus labrax*) genome (dicLab v1.0c) was selected as a reference and used for annotation procedures.

### RAD-Seq Mapping to the Reference Genome

Reads were mapped to the reference genome with a bowtie2 software package [32] using a set of global mapping parameters. DNA fragments were mapped along their entire length (from beginning to end). This approach has reduced the probability of non-orthologous mapping to relatively distant reference. After obtaining the *.sam files, we compress them to the *.bam format, sort and index the alignments, using the Samtools v1.7. SNP calling was performed using samtools and bcftools packages with maximum base quality - 30 (□□min□BQ parameter) [33].

In addition, the R packages: vcfR v1.8.0 [34], adegenet v2.1.1 [35], and ape v5.0 [36]were used for the subsequent genome analysis.

### Genotype clustering

We created a distance matrix for all samples, based on the dissimilarity of their genotypes and conducted a cluster analysis using the Neighbor-Joining method by applying the “nj” function in the ape 5.0 R-package, described by Paradis et al., (2019) [36] as a modern *ad hoc* phylogenetics analysis tool.

### Contamination test

In order to eliminate the suspicion of contamination, we determined the number of mismatches in the aligned data for each stickleback specimen. It is known that there is a number of “incorrectly” aligned sequences in any mapping data. These sequences, that show alternative nucleotides in alignment position, usually have low statistical support to characterize them as alternative alleles, because that algorithm identifies such deviations as mismatches. Moreover, DNA-library that has DNA contamination from another sample, should have a noticeably higher level of mismatches. Thus, the ratio of the number of mismatches to the total number of mapped nucleotides is an error rate, which is an indicator of contamination. To determine the error rate, we used samtools stats command [33], which estimated, besides other, mapping error rate, which is amount mismatches to total bases mapped according to cigar string information.

### Structure Software Analysis

The Structure software [37] input file containing the restriction site associated DNA (RAD) genotypes was created from *.vcf file using the PLINK v1.9 program [38]. The console version of the Structure program was compiled and launched on the NRC “Kurchatov Institute” computer cluster. The program was run several times with different parameters, but every time the same results were obtained. The publication included the results of the launch with the following parameters: 10,000 iterations of the burning period plus 20,000 Markov chain Monte Carlo (MCMC) replicas after burning. We used admixture ancestry and correlated allele frequency models for simulations. Uniform distribution of a priori parameters, without information about the sample origin population and geographical localization. The number of clusters – two.

### D-statistics with Admixtools Program Suite

The D-statistics method was used to formally evaluate whether a stickleback specimen displays DNA from a distantly related population. Indeed, the next three logical steps were applied to the admixture estimation:

1. *De novo* locus building from RAD-Seq data of stickleback specimens - requires assistance form the Stacks software package v.2.53 [39]. The denovo_map.pl pipeline was used for building stacks (loci) catalog and mapping reads from each specimen to the catalog.
2. The ancestor allele state was estimated to increase the test accuracy. Estimation was conducted only for derived alleles. We mapped each stack sequence to the reference (dicLab v1.0c, PRJEB5099) genome. The nucleotide, located in the SNP site on the reference genome, was considered an ancestor allele.
3. D-statistics estimation in Admixtools software suite [40] was used. Besides, the convertf and qpDstat tools were utilized for input file conversion and D-statistics and confidence values estimation.

### TreeMix Analysis

Graph-based models were used to determine the genomic admixture (migration) between different stickleback specimens; TreeMix package [41] was utilized in this analysis. The input file was converted from a genlight (adegent R package) object using the “gl2treemix” function of the dartR [42] R package. Before converting the TreeMix file, we created the multi-vcf file with the vcfR package, filtered the loci by genotyping quality “getQUAL” function for each locus - more than 500, and removed all loci, genotypes of which were the same for all samples. Tremix was launched with the parameter defining the number of migrations equal 2. The visualization of the ancestry graph and the migration was performed using the “plot_tree” R functions of the TreeMix package.

## Acknowledgments

This work has been carried out using computing resources of the federal joint usage center Complex for Simulation and Data Processing for Mega-science Facilities at NRC “Kurchatov Institute” (http://ckp.nrcki.ru/). Thanks to Prof. Azumi Aki for proof-reading of the final draft and Dr. Polina Nedoluzhko for ongoing support.

